# The translating bacterial ribosome at 1.55 Å resolution by open access cryo-EM

**DOI:** 10.1101/2022.08.30.505838

**Authors:** Simon A. Fromm, Kate M. O’Connor, Michael Purdy, Pramod R. Bhatt, Gary Loughran, John F. Atkins, Ahmad Jomaa, Simone Mattei

## Abstract

Our understanding of protein synthesis has been conceptualised around the structure and function of the bacterial ribosome^1–4^. This complex macromolecular machine is the target of important antimicrobial drugs^5^, an integral line of defence against infectious diseases. Here, we describe how open access to state-of-the-art cryogenic electron microscopy facilities combined with bespoke user support offered by the newly established EMBL Imaging Centre enabled structural determination of the translating ribosome from *Escherichia coli* at 1.55 Å resolution. The obtained structures allow for direct determination of the rRNA sequence to identify ribosome polymorphism sites in the *E. coli* strain used in this study and enables interpretation of the ribosomal active and peripheral sites at unprecedented resolution. This includes scarcely populated chimeric hybrid states of the ribosome engaged in several tRNA translocation steps resolved at ~2 Å resolution. The current map not only improves our understanding of protein synthesis but also allows for more precise structure-based drug design of antibiotics to tackle rising bacterial resistance.

## Main

Cryogenic electron microscopy (cryo-EM) allows for detailed structural determination of biomolecules, and it is currently the method of choice for obtaining high-resolution reconstructions of large and conformationally heterogenous macromolecular complexes vitrified in functional buffers after purification or directly within their native cellular context^6–8^. Although a remarkably powerful tool for structural biologists, cryo-EM experiments still rely on access to both high-end instrumentation and advanced expertise in sample preparation, data acquisition and image processing, thus posing major challenges for a growing number of life scientists that approach this method to tackle their biological questions for the first time. To address the ever-increasing demand for access to cryo-EM infrastructure and expertise, several local and national facilities have been established to provide access to cutting-edge transmission electron microscopes and project support by expert staff. To this end, the European Molecular Biology Laboratory (EMBL) has recently established the EMBL Imaging Centre, a new service unit with the main mission to rapidly make the most advanced microscopy technologies available as a synergistic service portfolio to the international user community while providing comprehensive support from sample preparation to image acquisition and data analysis.

To demonstrate the capabilities of our cutting-edge instrumentation and experienced team to support external user projects requiring cryo-EM single particle analysis workflows, we vitrified, imaged, and analysed purified *E. coli* ribosomes stalled during translation. The bacterial 70S ribosome is an ideal candidate due to its fundamental importance in protein synthesis as well as being an active antibiotic target for antimicrobial research. Furthermore, capturing the ribosomes intrinsic conformational states, that usually represent challenges for structural biology approaches, is imperative to gain an atomic level understanding of the function of this intricate machine.

The ribosome has been a suitable target for single particle cryo-EM due to its large size, globular shape, and high RNA content, which generates relatively high contrast in the otherwise noisy cryo-EM images. Despite this, high-resolution structures (< 2 Å resolution) reported recently were limited to protein complexes with high symmetry such as apoferritin^9–12^, aldolase^12^ or β-galactosidase^13^. In contrast, the most detailed reconstructions of ribosomal complexes were so far resolved to 2.0 Å and 2.2 Å resolution, for the bacterial and the mammalian ribosomes^14,15^, respectively. This was mostly due to the lack of intrinsic symmetry of the ribosomes and their conformational and compositional heterogeneity, deriving from inter-subunit movements, flexible expansion rRNA segments, and the intermittent interactions with a plethora of additional translational factors involved in the different steps of translation.

Here, we describe how open access to state-of-the-art transmission electron microscopes combined with bespoke user support enabled structural determination of the bacterial translating ribosome at 1.55 Å resolution. The structure resolved many important features of the ribosome at atomic detail including chemical modifications, magnesium, and potassium ions, as well as water molecules. The current resolution also allowed for direct identification of multiple rRNA variations including a base pair swap in one of the rRNA helices that is distinct between different *E. coli* strains. The cryo-EM structure contained highly resolved features in the periphery of the ribosome including the ribosomal protein uL9, which provides insights on its possible function. Finally, the current dataset allowed us to resolve several conformational states of the ribosome with different tRNAs from a relatively small number of particles (~20,000 particles) at ~ 2 Å resolution.

## Results

### Isolation of translating ribosomes for cryo-EM structure determination

To isolate bacterial ribosomes actively engaged in translation, we employed a cell free *in vitro* translation PURE system^16,17^ of a mRNA encoding a FLAG-tagged nascent chain used for purification. The reaction mixture was incubated at 37°C to allow the ribosome to engage mRNA and start several elongation cycles, and after 20 minutes was placed on ice to pause translation. The translating ribosomes were isolated via a FLAG-tag affinity purification step followed by an ultracentrifugation step to pellet ribosomes from the preparation. Purified ribosomes were then applied to cryo-EM grids for imaging. We rationalised that the sample would contain translating ribosomes frozen at different functional states that can be further purified *in silico* by a combination of 3D image classification and focused 3D refinements approaches.

### Cryo-EM sample screening at the EMBL Imaging Centre

To produce and identify a cryo-EM grid suitable for high-end data collection on a Titan Krios microscope, a total of 19 grids were vitrified over three freezing sessions and screened in two sessions using a Glacios (ThermoScientific) transmission electron microscope. At first, the screening was aimed at optimising sample spreading on the grid and ice thickness. Here, two different vitrification devices (ThermoScientific Vitrobot Mark IV and Leica EM GP2) and multiple different grid types (holey carbon grids with and without an additional layer of continuous carbon) were employed. Grid quality assessment was based on the low magnification grid overview which was acquired for each grid (commonly termed ‘grid map’ or ‘grid atlas’) and high magnification screening images from those grids which showed promising ice thickness and had sufficient imageable areas for potential high-resolution data collection (Fig. 1 and Extended Data Fig. 1).

**Fig. 1:**
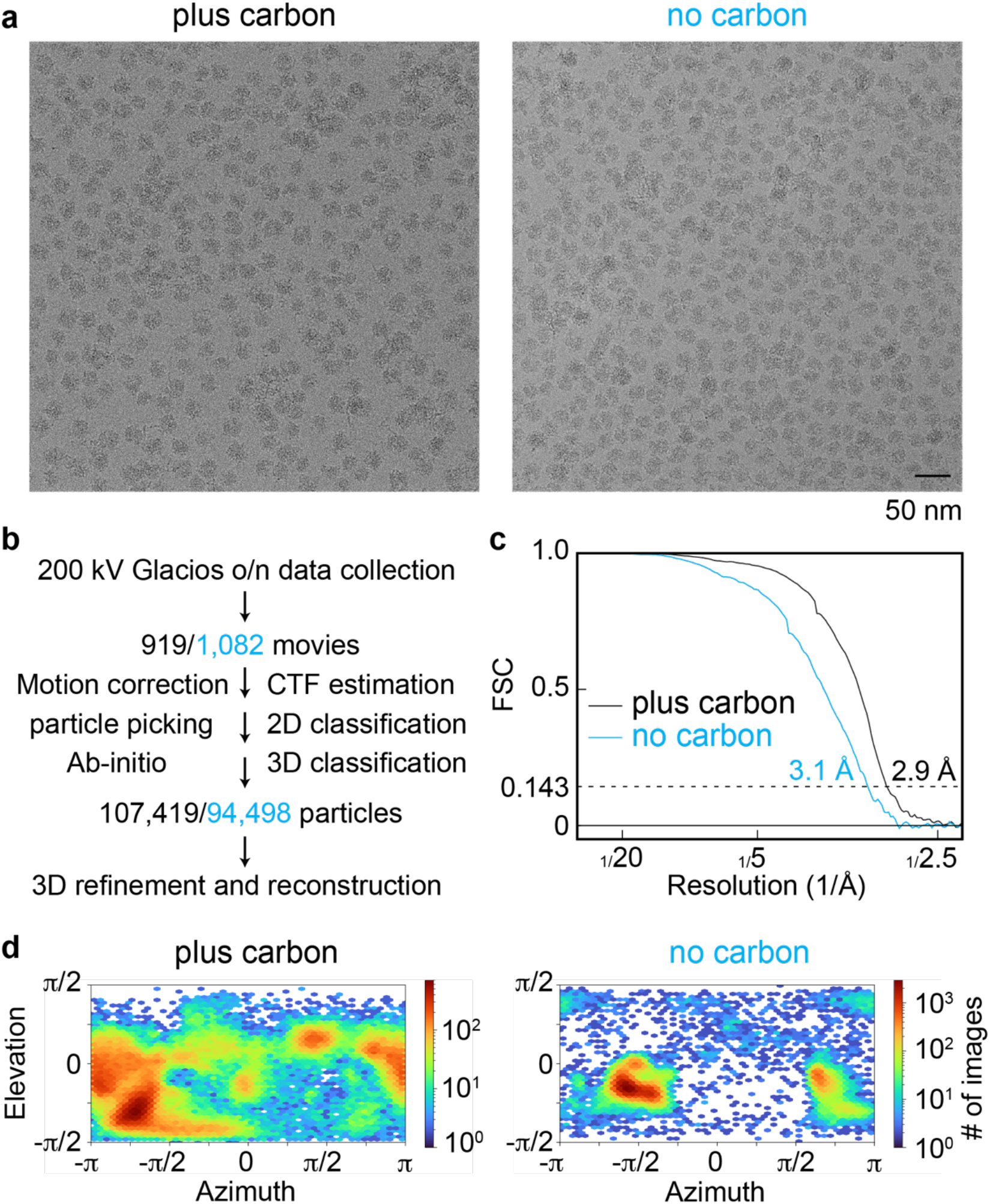
Initial Cryo-EM sample screening to select the grid for high-resolution data collection. **a**, Cryo-EM micrographs of the ribosome particle images applied to holey carbon grids with (left) and without (right) an additional thin layer of continuous carbon applied (2 nm). **b**, Sample screening processing scheme using cryo-EM data collected on ThermoScientific Glacios electron microscope. **c**, Fourier Shell Correlation (FSC) plots for the data collected on holey grids with and without an additional layer of continuous carbon applied. **d**, Orientation distribution of the particles from data collected on plus and no carbon grids.

During the second stage of screening, the selected grids were assessed for ribosome particle density and distribution. A small overnight data set (~1,000 movies) was then collected from the most promising grid at the end of each Glacios screening session. The two selected grids were a Quantifoil R2/1 grid with an additional 2 nm layer of continuous carbon vitrified with the Vitrobot (‘plus carbon’ grid) and a Quantifoil R2/1 grid without the additional layer of continuous carbon vitrified with the EM GP2 using back side blotting to increase particle density in the foil holes (‘no carbon’ grid) (Fig. 1a). Each dataset was analysed by employing the same processing workflow yielding 3D reconstructions of the 70S ribosome at ~3 Å resolution from approximately 100,000 particles (Fig. 1b and c). Despite the similar nominal resolutions obtained, the ‘plus carbon’ dataset had a substantially better particle orientation distribution compared to the ‘no carbon’ data set which suffered from severe preferred orientation (Fig. 1d). Therefore, we selected the ‘plus carbon’ grid for further high-end data collection on a Titan Krios G4 (ThermoScientific) equipped with a cold-FEG electron source, Selectris X energy filter and Falcon4 direct electron detector; over a 3-day session, a total of ~20,000 movies were collected.

### High-resolution structure of P-site bound translating ribosome

We chose to process particles using a combination of both cryoSPARC^18^ and RELION^19^ software. Particle picking was done semi-automatically with crYOLO using the pre-trained general model^20^, then followed by standard 2D classification and *ab initio* model generation which suggested the sample contained both translating ribosomes and free 50S subunit particles. Heterogeneous refinements in cryoSPARC were then employed to remove particles that belong to the free large ribosomal subunit and resulted in ~750,000 70S ribosome particles. Further 3D classification resolved a class containing ~650,000 particles of a translating ribosome containing P-site tRNA and a weaker density for the E-site tRNA with the small 30S subunit in the unrotated state (Extended Data Fig. 2-4). This is a widely observed state for translating ribosomes isolated using *in vitro* translation systems^21–23^. In this map, the P-site tRNA is tethered to a nascent chain (NC) density with weak density traceable to the backbone of the alpha carbons (Extended Data Fig. 5). However, clear densities of side chains were not observed in the polypeptide exit tunnel due to averaging of several states during translation and thus we did not fit an atomic model in this area.

A 3D variability analysis focused on the tRNA sites identified two subclasses with P- and E-site (~500,000 particles) and A, P and E-site (~150,000 particles) tRNA densities, respectively. A combination of 3D refinement schemes, CTF refinement schemes as well as particle polishing^19^ were then used to obtain a final reconstruction to 1.55 Å overall resolution of the larger class containing EM-densities corresponding to P- and E-site tRNAs (Extended Data Fig. 2 and 4, Extended Data Table 1). The local resolution of the small subunit was slightly lower (~1.9 Å) and was further improved to ~1.7 Å after performing focused 3D refinements and signal subtraction of the large subunit (Extended Data Fig. 4).

In the obtained structure, we can visualise rRNA modifications and protein side-chain conformers (Fig. 2) similar to those observed in a previously reported structures of the bacterial ribosome bound to mRNA and tRNA at 2.0 Å^15^. At the current resolution, we can observe separate EM-densities of water molecules coordinating magnesium ions bound to the RNA backbone (Fig. 2b). In addition, we resolved EM-densities for potassium ions (Fig. 1b), based on their identified positions in a recent x-ray structure^24^. Although the state of the ribosome is paused at different states during translation, we visualised the mRNA with defined anticodon-codon interactions being visible (Extended Data Fig. 6). It is thus likely that the ribosome is paused at preferred sites due to visible densities of purine and pyrimidine rings. However, a reliable assignment of the pausing site could not be assigned and is not further discussed here.

**Fig. 2:**
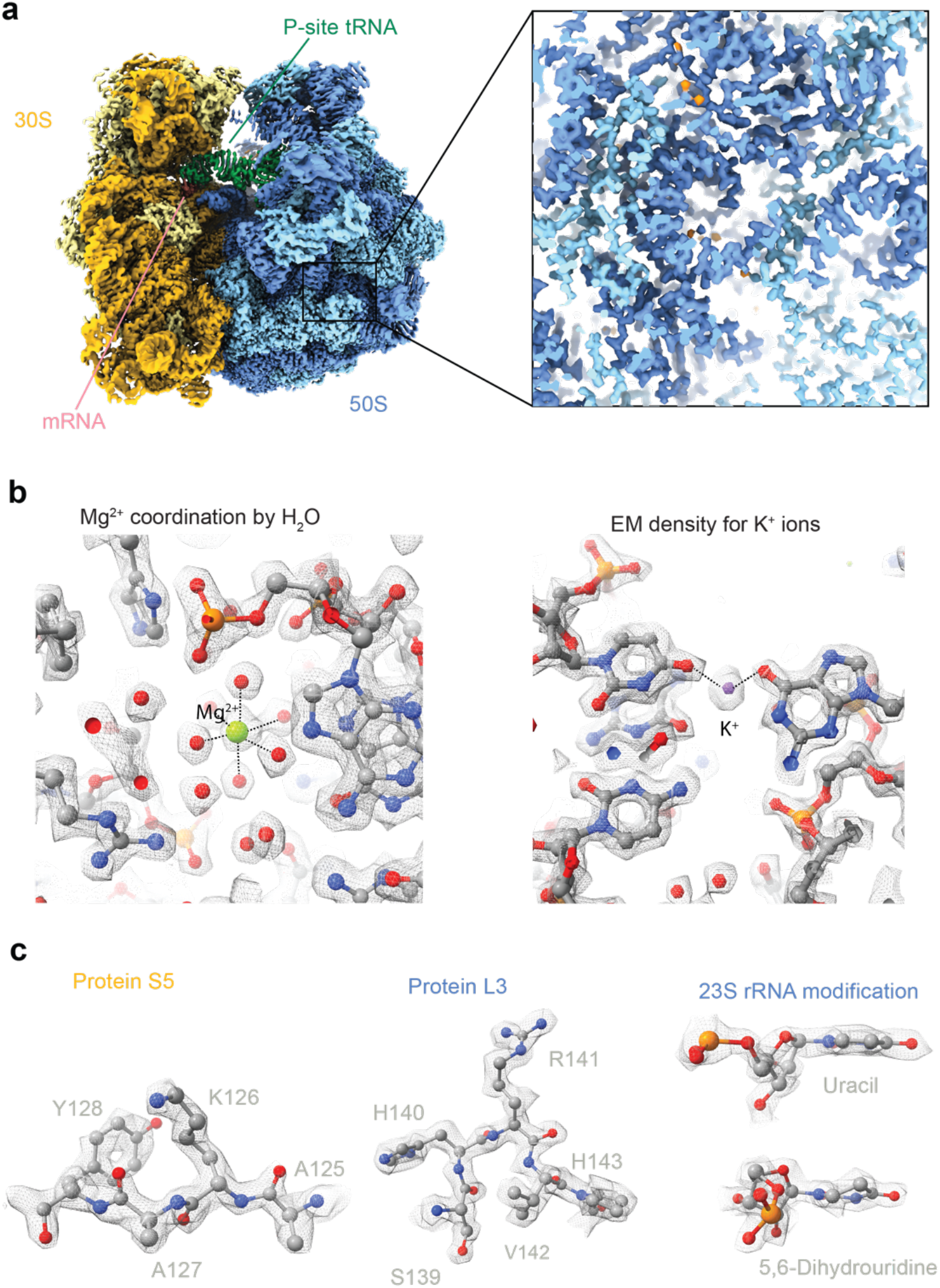
1.55 Å resolution reconstruction of the 70S ribosome. **a**, Cryo-EM map of the bacterial translating ribosome at 1.55 Å coloured in dark yellow (16S rRNA), light yellow (ribosomal proteins in the small subunit), light blue (large subunit ribosomal proteins), dark blue (23S and 5S rRNA). **b** and **c**, Closeup densities of protein side chain conformation, water molecules, magnesium, and rRNA modifications are shown as sticks with overlaid EM-densities shown as mesh.

### A C-G base-pair swap in the 23S rRNA observed in the high-resolution map

Interestingly, we observed a mismatch between the *E. coli* MRE600 strain sequence of the 23S rRNA docked into our cryo-EM map. The initially identified mismatch was a G-C base-pair swap in the position G2209 and C2215 in helix 79 of the 23S rRNA. After closer inspection, we realised that the PURE *in vitro* translation system kit was developed based on the *E. coli* B-strain instead^25,26^, of which the sequence fits well into the EM-density of our maps. Through sequence alignments of the seven different 23S rRNA operons of the *E. coli* B-strain and comparing their fit to our EM density we could identify loci C2566_01480 and C2566_04790 as the predominant ones in our 70S ribosome sample. There are two additional mutations within the same rRNA helix which fit to our density (Fig. 3a and b). All other differences between the MRE600 and B strain rRNA sequences (mutations and insertions), are also reflected by our EM map. Sequence polymorphisms within the 23S and 16S rRNA is a common property of bacterial strains and environmental isolates of microbes which could be used as a possible indicator of strain virulence^27,28^. The C-G mismatch identified here is an intriguing example of co-evolution where two separate mutations in the 23S rRNA sequence were attained to maintain the base-pairing and the integrity in the 23S rRNA helix.

**Fig. 3:**
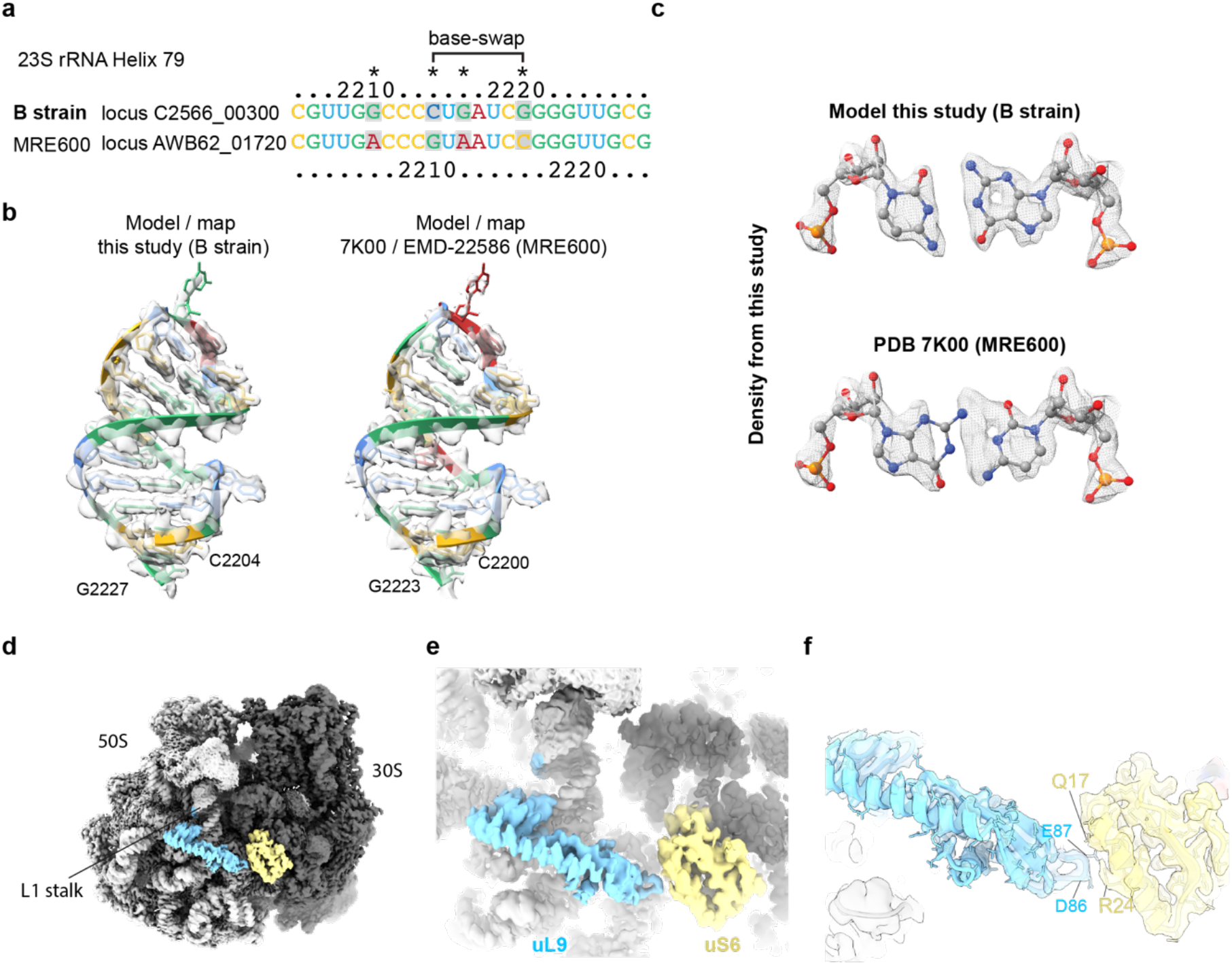
Direct Sequencing of rRNA polymorphism and full-length UL9 resolved by the high-resolution 70S structure. **a**, Partial sequence alignment of H79 of the 23S rRNA between the *E. coli* B-strain and MRE600. Base-swap positioning is indicated. Other observed mutations are indicated by an asterisk. **b**, Snapshot of H79 shown as cartoon illustration with base colouring as in “a”. Overlaid cryo-EM density is shown as transparent isosurface. **c**, Close-up of the base pair swap using B-strain sequence and MRE600 sequence (PDB 7K00) with overlaid EM density shown as mesh. **d**, uL9 (cyan) and uS6 (yellow) highlighted in the context of the cryo-EM map of the 70S ribosome. Large subunit is in light grey, small subunit in dark grey. L1 stalk of the ribosome is indicated. **e** and **f**, Closeups of uL9 density with surrounding interactions with the small subunit. Ribosomal protein uS6 is indicated. Overlaid EM density is shown as a transparent surface in panel f. Colouring as in panel d.

### Density corresponding to the ribosome protein uL9

In the current map, the spread of local resolution at the periphery is around 2-3 Å (Extended data fig. 4), which underscores the stability of our sample preparation and data processing schemes. We observe a well-resolved density at side-chain resolution of the ribosome protein uL9 (Fig. 4d-f). uL9 is located at a peripheral position of the ribosome forming a strut-like shape below the L1 stalk and bridging the large and the small subunits of the ribosome. In the current structure, uL9 is observed establishing contacts with uS6 where it locks the L1 stalk in the closed conformation contacting the E-site tRNA (Fig 4d-e). The current conformation of this protein is stabilised via a salt bridge between Arg24 (S6) and Glu87 (uL9) (Fig. 4f). Notably, the strut-like shape of uL9 is rich in negatively charged residues through its extended structure (Extended Data Fig. 7), which would otherwise repel away from the ribosome. We speculate that this unusual feature of a ribosomal protein carries an important function during protein synthesis that could also contribute to its detachment and flexibility in previous x-ray and cryo-EM structures that were done under variable salt conditions.

**Fig. 4:**
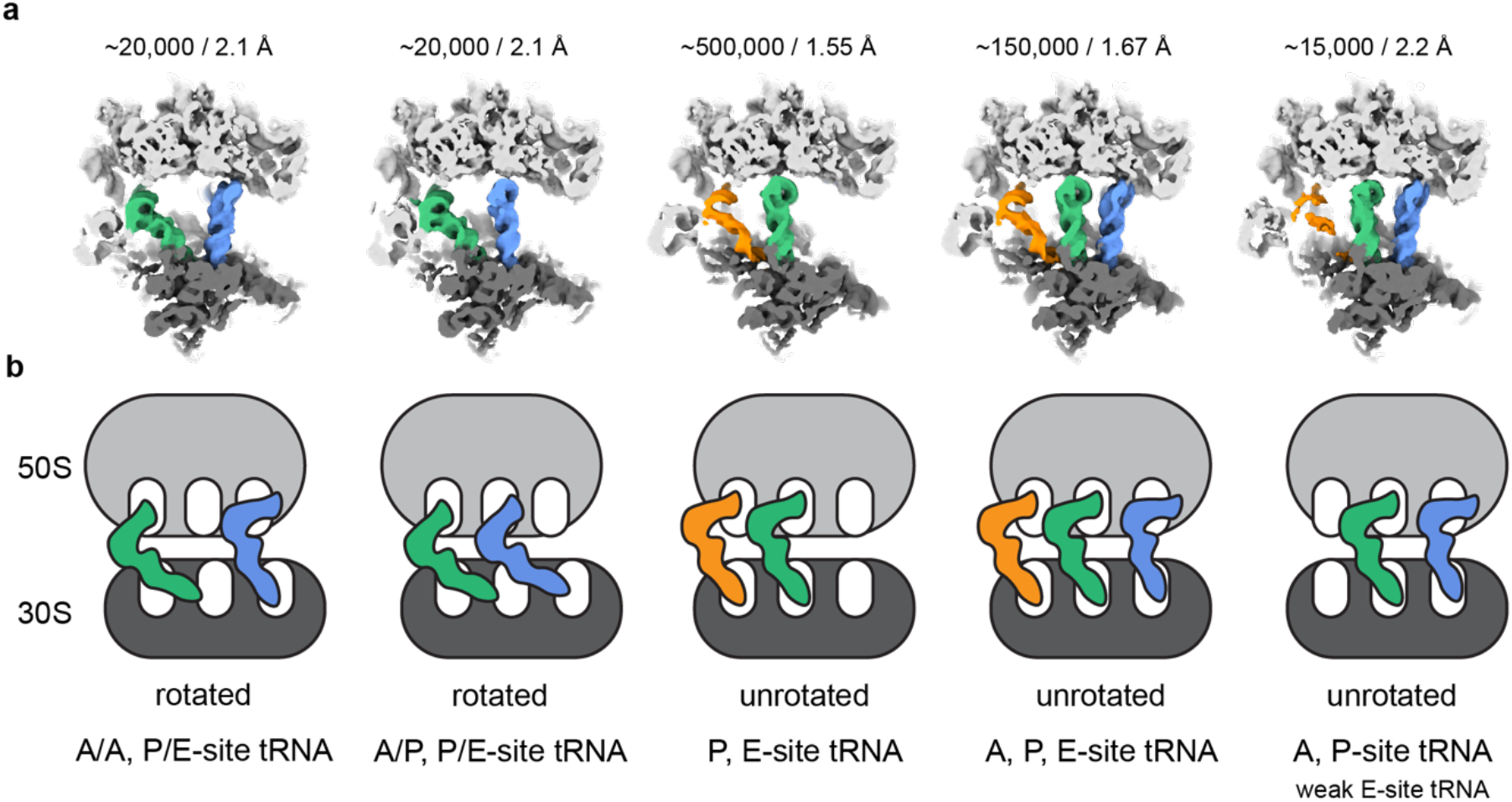
Different tRNA states revealed by 3D classification and variability analysis at high resolution from small number of particles. **a**, Cryo-EM densities of different ribosome translating states from 3D variability analysis low-pass filtered at 5 Å. Large ribosomal subunit coloured in light grey, small subunit in dark grey, A-site tRNA in blue, P-site in green, E-site in orange. Number of particles and resolution of the 3D reconstruction for each class are indicated above. **b**, Schematic representation of all identified tRNA states in the data set. Colouring as in “a”. The respective tRNA state is indicated below.

The ribosomal protein uL9 plays an important role in influencing reading frame maintenance and restraining forward slippage on mRNA. This restraint can be circumvented by recoding signals that cause programmed frameshifting that are availed of by many viruses and mobile elements^29–31^. Recently, L9 interactions across neighbouring ribosomes translating the same mRNA were also shown to play a role in ribosome quality control pathways bridging interaction between two colliding ribosomes^32,33^. High-resolution of this protein is thus highly pertinent to determining the outstanding functional aspects of this enigmatic dual-mode protein.

### Multiple functional states of the translating bacterial ribosome

We also investigated the presence of additional functional states of the translating ribosome in the dataset. We employed 3D classification to identify a class of ribosomes in the rotated state which was further sub-classified using the 3D variability approach implemented in cryoSPARC by focusing on the tRNA binding sites on the ribosome (Fig. 4, Extended Data Fig. 2 and 3). This classification scheme resolved three tRNA translocation intermediates of the translating ribosomes. Besides the above described classical unrotated state with bound A-, P-, and E-site tRNAs, we also resolved two additional states containing chimeric state rotated ribosomes. The first class is of A/P- and P/E-tRNA hybrid state ribosomes and the second class with A- and P/E-site tRNAs (Fig. 4). The chimeric state was observed in the absence of the tRNA-bound elongation factor EF-Tu and was described in more detail in recent cryo-EM structures^34–36^. Interestingly, each of the two intermediates represent only ~2.7 % (~20, 000 particles) of the total dataset and yet could be refined to ~2.1 Å indicating the possibility to sort for rare intermediates that can be resolved to high resolution from a relatively small number of particles.

## Discussion

In summary, we present multiple structures of the *E. coli* translating ribosome, with the most populated state resolved at a global resolution of 1.55 Å. To our knowledge, this is currently the highest resolution reported for the ribosome or any non-symmetrical complex by single particle cryo-EM. Determining structures of the ribosome has continuously progressed over the last 25 years since the initial x-ray structures emerged^37–41^. More recently, cryo-EM has provided insight into a plethora of functional states describing how ribosomes function, some of which were de-convoluted from the same reaction at high resolution. With current developments in cryo-EM image processing, direct pull-downs of active complexes are becoming the go-to approach requirement for resolving short-lived intermediates.

We suggest that a combination of improvements in sample preparation, microscopy hardware, data collection, and processing schemes with respect to previous studies allowed for this substantial advancement of the achievable resolution for an asymmetric and compositionally heterogeneous complex like the ribosome. The highly resolved rRNA in the presented structure allowed for sequence identification of portions of the rRNA directly from the map, where we detected a C-G base pair swap in H79 of the large subunit rRNA. This swap is due to sequence variation in the *E. coli* B strain, compared to the MRE600 strain. Interestingly, despite the mutations being in two distant positions, the base pair swap does not alter the structural integrity of the helix. Such possible identification of base pair swapping in the rRNA could be used to identify bacterial strains by directly performing sequence analysis on their rRNA using cryo-EM performed on translating ribosomes purified from complex bacterial populations. Finally, the current dataset demonstrates how the latest generation of microscopy hardware and software allowed for the 3D classification and refinement of multiple conformational states of asymmetric complexes from a relatively small number of particles (~ 20,000) to around ~ 2Å. The results presented here show the importance of imaging facilities that provide open access to state-of-the-art instrumentation and tailored support to enable advanced structural studies pursued by researchers approaching cryo-EM as a new method.

## Methods

### Sample Preparation

A synthetic DNA sequence (gBlock, IDT) with a T7 promoter, a leader region, and a Shine Dalgarno sequence upstream of sequence encoding an N-terminal 3x FLAG-tag followed by a 96 aa linker and sequence (373 nt) from bacteriophage MGS13 was used as template for PCR using primers gBlock S and gBlock AS (see Extended Data Table 2). PCR amplicons were verified by agarose gel electrophoresis and purified using the Monarch PCR and DNA Cleanup Kit (New England Biolabs). *In vitro* transcription was performed using the T7 RiboMAX Express Large Scale RNA Production System (Promega), followed by Lithium Chloride precipitation to purify the synthesized RNA.

In vitro translation was carried out using the deltaRF123 Kit (New England Biolabs). 200 ng of mRNA was added per μL of reaction. Reactions were incubated at 37 °C for 30 minutes. Translation reactions were chilled on ice for 10 minutes to halt translation, and HEPES KOH pH 7.4 was added to a final concentration of 50 mM. Chilled translation reactions were directly incubated with anti-FLAG M2 beads (Sigma) for 2 hours at 4°C with gentle mixing.

The beads were then washed with 50 mM HEPES, pH 7.4, 100 mM KOAc, 25 mM Mg(OAc)2, 0.1% Triton X-100, 1 mM DTT; 50 mM HEPES, pH 7.4, 250 mM KOAc, 25 mM Mg(OAc)2, 0.5% Triton X-100, 1 mM DTT and RNC buffer (50mM HEPES, pH 7.4, 100 mM KOAc, 25 mM Mg(Oac)2, 1mM DTT). RNCs were eluted after 3 sequential 10 min incubations at room temperature in the RNC buffer that contained 0.2 mg/mL 3x FLAG peptide (Sigma). The elutions were combined and centrifuged at 186,000 x g at 4 °C for 1.5 h in a (TLA-120.1) rotor (Beckman Coulter). Supernatant was discarded and the pellet was resuspended in the RNC buffer at a concentration of ~200 nM. At each step of translation reactions and purification, aliquots were taken to perform SDS-PAGE on 4-12% Bolt Polyacrylamide gels (Invitrogen) and Western blots. Gene products of interest were probed using a commercial anti-FLAG antibody (Sigma Cat. No. F1804).

### Cryo-EM sample preparation

Cryo-EM grids have been prepared with a Vitrobot Mark IV (plus carbon sample, Thermo Scientific) or an EM GP2 (no carbon sample, Leica) plunger. Environmental chambers were set to 6 °C and 100 % humidity. Grids were rendered hydrophilic by plasma cleaning for 30 s in a 90/10 % Ar/O mixture with a Fischione 1070 plasma cleaner right before plunge freezing. 2.5 μl of the undiluted ribosome sample were applied to the foil side of the grid. The plus carbon sample (Quantifoil Cu300 R2/1 + 2 nm C) was blotted for 2 s with blot force +3 after a 30 s wait time after sample application and immediately plunge frozen in liquid ethane. The no carbon sample (Quantifoil Cu200 R2/1) was blotted for 1.5 s from the back side right after sample application and immediately plunge frozen in liquid ethane. In all cases Whatman 597 blotting paper was used.

### Cryo-EM Data Collection

Grid screening and screening data acquisition was performed on a Glacios TEM operated at 200 kV equipped with a Selectris X energy filter and Falcon 4 direct electron detector (ThermoScientific). Data was acquired at a nominal magnification of 100 kx resulting in a calibrated pixel size of 1.154 Å on the camera. A 50 μm C2 and a 100 μm objective aperture were inserted and the width of the energy filter slit was set to 10 eV. The TEM was operated in nanoprobe mode at spot size 6. Movies were acquired over 8.3 s exposure time with a total accumulated dose of ~40 e^-^/Å2 (~6.4 e/px/s over an empty area on the camera level) in counting mode and saved in the EER file format. For the plus carbon data set, 919 movies were acquired in one overnight data collection (1 exposure per hole/stage movement) and for the no carbon data set 1,082 movies were acquired in the same way.

After evaluation of the two screening data sets, the plus carbon grid was selected for high-resolution data acquisition on a Titan Krios TEM operated at 300 kV equipped with a C-FEG, Selectris X energy filter and Falcon 4 direct electron detector (ThermoScientific). A total of 19,449 movies were collected over three days at a nominal magnification of 165 kx resulting in a calibrated pixel size of 0.731 Å on the camera. A 50 μm C2 and a 100 μm objective aperture were inserted and the slit width of the energy filter was set to 10 eV. The TEM was operated in nanoprobe mode at spot size 5 and a beam diameter of ~460 nm. Movies were acquired over a 5.2 s exposure time with a total accumulated dose of ~40 e^-^/Å^2^ (~4.1 e/px/s over an empty area on the camera level) in counting mode and saved in the EER file format (1,293 fractions per movie). The target defocus was set to −0.5 to −1.5 μm with 0.1 μm steps between holes. 16 movies were acquired per hole using beam image shift (11 in the outer ring, 5 in the inner ring) before moving to the next hole by stage shift. All TEM screening and data acquisition was performed using SerialEM^42^.

### Data Processing and Model Building

CryoEM data processing workflows for the Glacios screening data sets and the Krios high-resolution data set are shown in Fig. 1 and Fig. S2-3, respectively. In brief, motion correction of EER movies was performed with the CPU implementation of MotionCor2 from Relion4^43^. Initial CTF estimation was performed using CTFFIND4^44^. Gautomatch (http://www.mrc-lmb.cam.ac.uk/kzhang/) was used for particle picking of the Glacios screening data, crYOLO^20^ with the pre-trained general model was used for picking of the high-resolution Krios data set. Particle extraction, 3D classification with alignment as well as Bayesian polishing were performed with Relion4. All other cryoEM data processing steps were performed in cryoSPARC^18^. Separation of 50S and 70S particles was achieved via Heterogeneous Refinement, different tRNA states were identified using a combination of 3D classification and 3D Variability analysis. Maps were post-processed by local filtering after local resolution estimation as implemented in cryoSPARC. Conversion of cryoSPARC .cs files to Relion .star files was performed using UCSF pyem^45^ in combination with in-house written bash scripts (https://github.com/simonfromm/miscEM). All final Homogeneous Refinements were done with per-particle defocus and per-group CTF parameter optimization as well as Ewald Sphere Correction switched on. Exposures from all beam image shift positions of the acquisition pattern were put into a separate group resulting in a total of 16 exposure/optics groups.

Model building was started with individually rigid-body fitting all chains from the published 2 Å *E. coli* 70S ribosome structure (PDB 7K00) in COOT^46^ After manual inspection of the rRNA of the large and small ribosomal subunits for mutations, the residue numbering of the *E. coli* B-strain was used. All chains were inspected and adjusted manually in COOT and subsequently refined with real-space refinement in PHENIX^47^ against the unsharpened map with secondary structure and Ramachandran restraints switched off (Extended Data Table S1). The C-terminal part of uL9 absent from PDB 7K00 was built de novo based on the AlphaFold2 model. Water molecules were added to the 30S and 50S part of the structure separately using ‘Find Waters’ in COOT. Mg and K positions were inferred from PDB 7K00 and 6QNR, respectively. All Mg and K ion positions were checked and modified manually when necessary. The model geometry was validated using MolProbity^48^. The model vs. map FSC at FSC=0.5 coincides well with the one determined between the map half-sets at FSC=0.143 (Extended Data Fig. 4). All cryoEM density and model renderings were generated with ChimeraX^49^.

## Data availability

The maps and models are deposited in the EM and protein database EMD-XXXX, EMD-YYYY, and EMD-ZZZZ and PDB-XXX. All micrographs and coordinates for the picked particles are deposited in Electron Microscopy Public Image Archive as EMPIAR-XXXX.

## Author contributions

S.M., A.J. and J.F.A. conceived the project, K.O., P.R.B., G.L. purified the ribosomes. S.A.F. screened and collected cryo-EM data. S.A.F., A.J., and M.P. processed the cryo-EM data. S.A.F built the model with input from A.J. and S.M.. S.M., A.J. and J.F.A. supervised the work. A.J., S.A.F., and S.M. wrote the manuscript. All authors contributed to the final version of the manuscript.

## Acknowledgements

We would like to thank members of the Mattei, Jomaa and Atkins labs for discussions. We are grateful to Dr. Jochen Zimmer for his comments on the manuscript. We also thank Maciej Gluc for discussions. We acknowledge the access and services provided by the Imaging Centre at the European Molecular Biology Laboratory (EMBL IC), generously supported by the Boehringer Ingelheim Foundation.

## Extended data figures and tables

**Extended Data Fig. 1.**
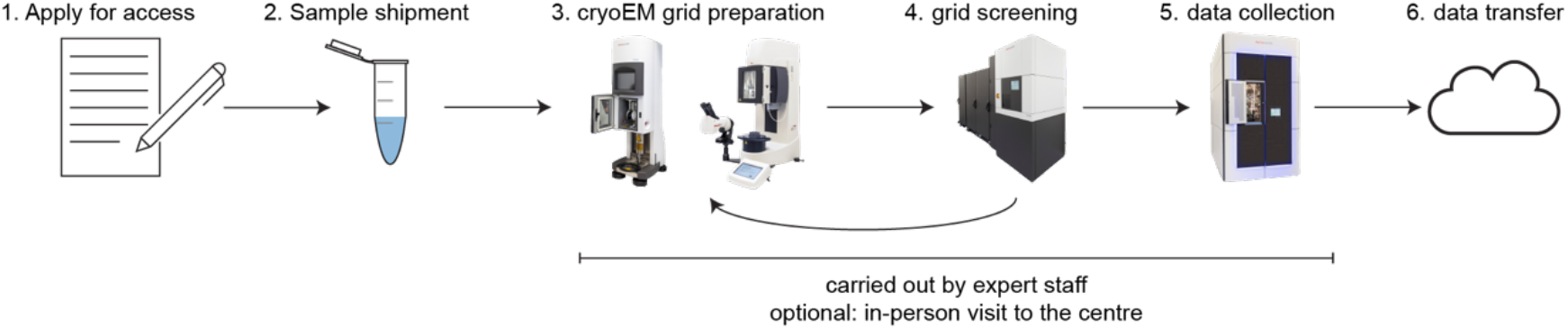
General access scheme/workflow. A schematic of the workflow for cryo-EM sample preparation and data collection at the EMBL Imaging Centre.

**Extended Data Fig. 2.**
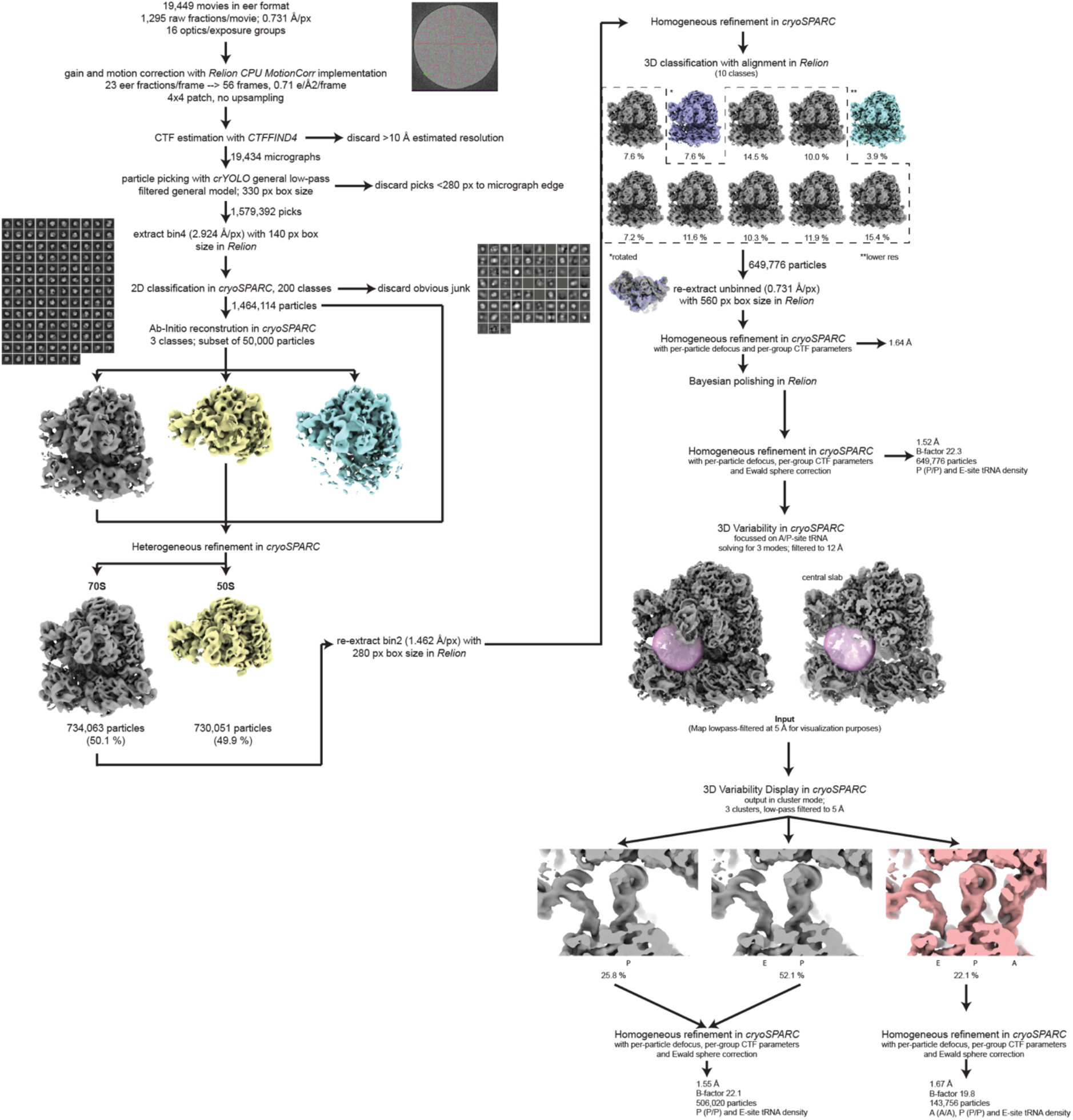
Processing scheme of the ribosome in the unrotated state. Processing scheme to initially separate 70S ribosomes from free 50S subunits. 70S ribosomes were then classified using heterogenous refinement in cryoSPARC, which separated ribosomes based on the rotated and unrotated states of the small ribosomal subunit. Ribosomes in the unrotated state were then classified using a binary mask around the tRNA binding site to resolve additional states of bound tRNA. 50S subunit were not further processed.

**Extended Data Fig. 3.**
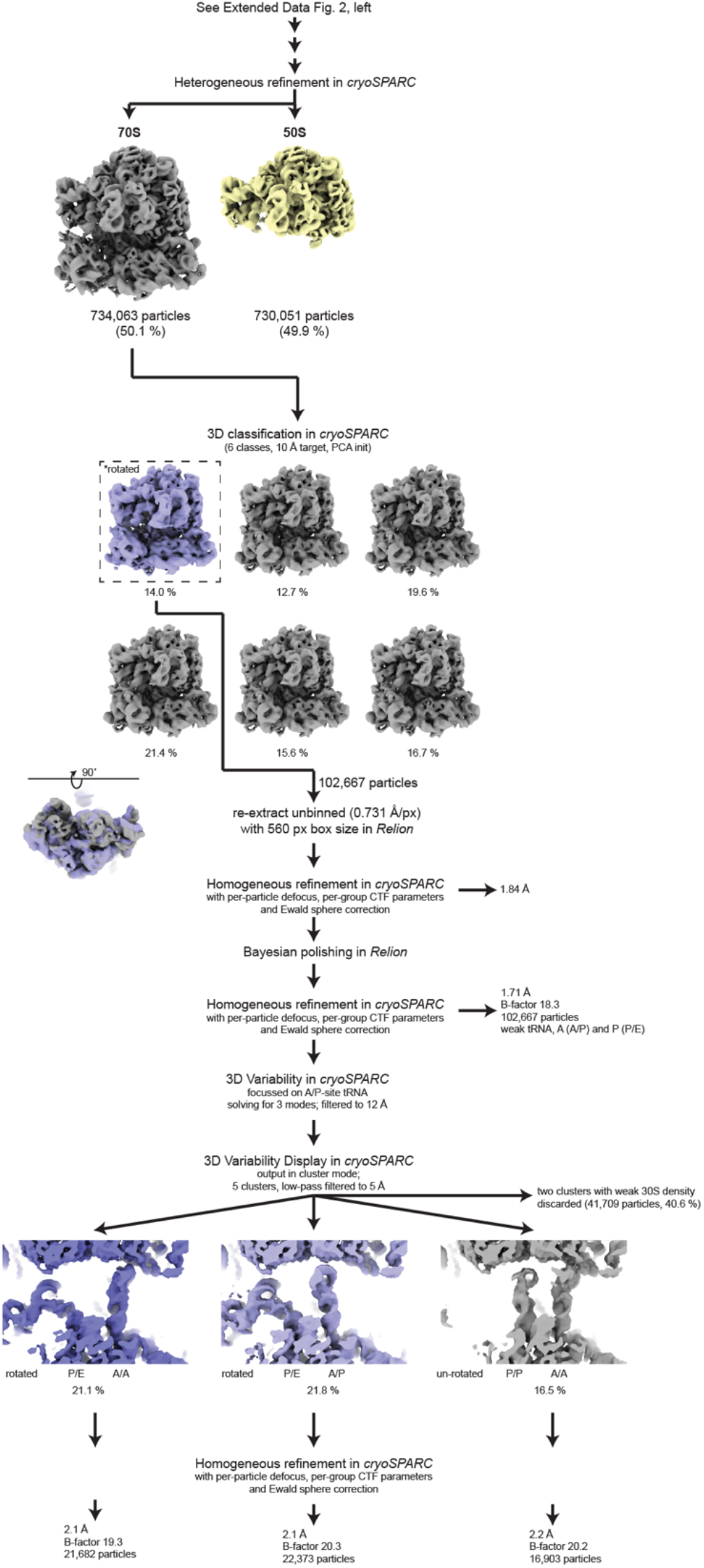
Processing scheme for the ribosome in the rotated state. Ribosomes in the rotated state were classified using a binary mask around the tRNA binding site to resolve additional states of bound tRNA, similarly as described in Extended Data Fig 2.

**Extended Data Fig. 4.**
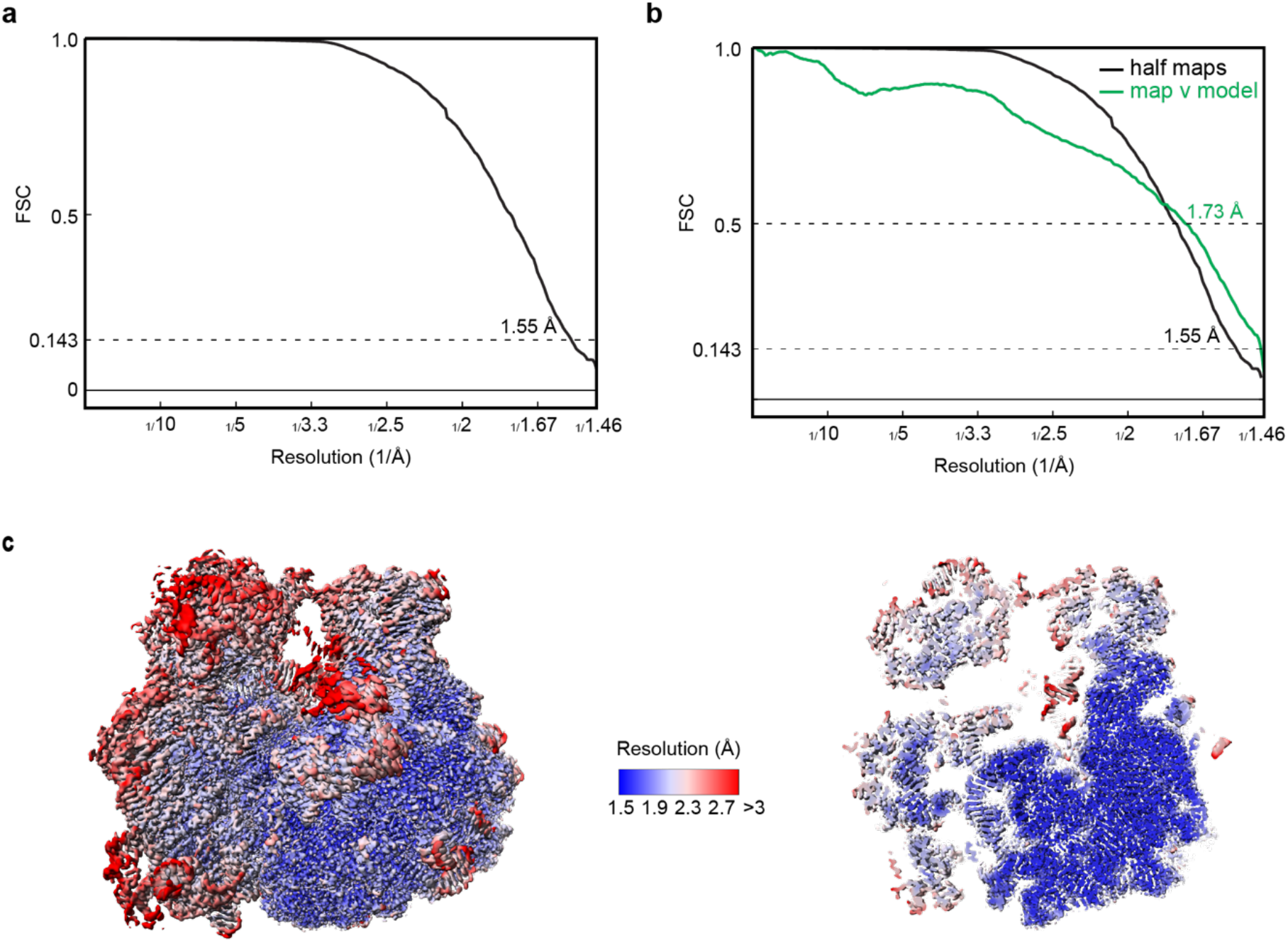
FSC and Local resolution plot of the high-resolution E. coli translating ribosome. **a**, Gold-standard FSC of final 3D reconstruction **b**, Map-model validation: gold-standard FSC (black) overlaid with map-vs-model FSC (green). **c**, Local resolution estimation. Left, surface representation of the full ribosome. Right, central slice through the density.

**Extended Data Fig. 5.**
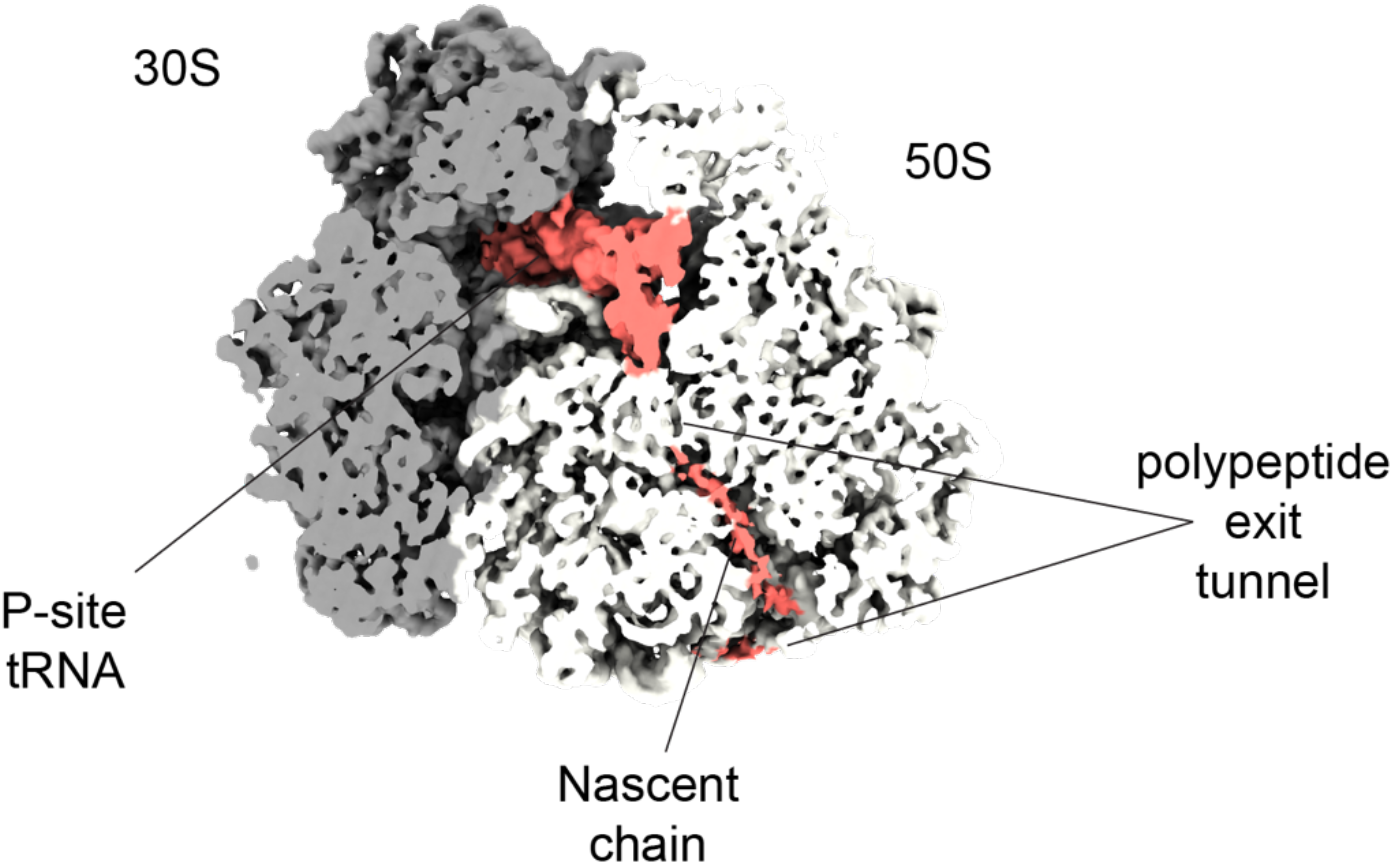
Visualisation of the tRNA-tethered nascent chain in the ribosomal polypeptide exit tunnel. Cross-section of the polypeptide exit tunnel with resolved EM-density for the nascent chain. Ribosomal subunits 30S and 60S are coloured light and dark grey to emphasise the density of the nascent chain shown in red. Cryo-EM density is shown as surface and low-pass filtered to 5 Å resolution.

**Extended Data Fig. 6.**
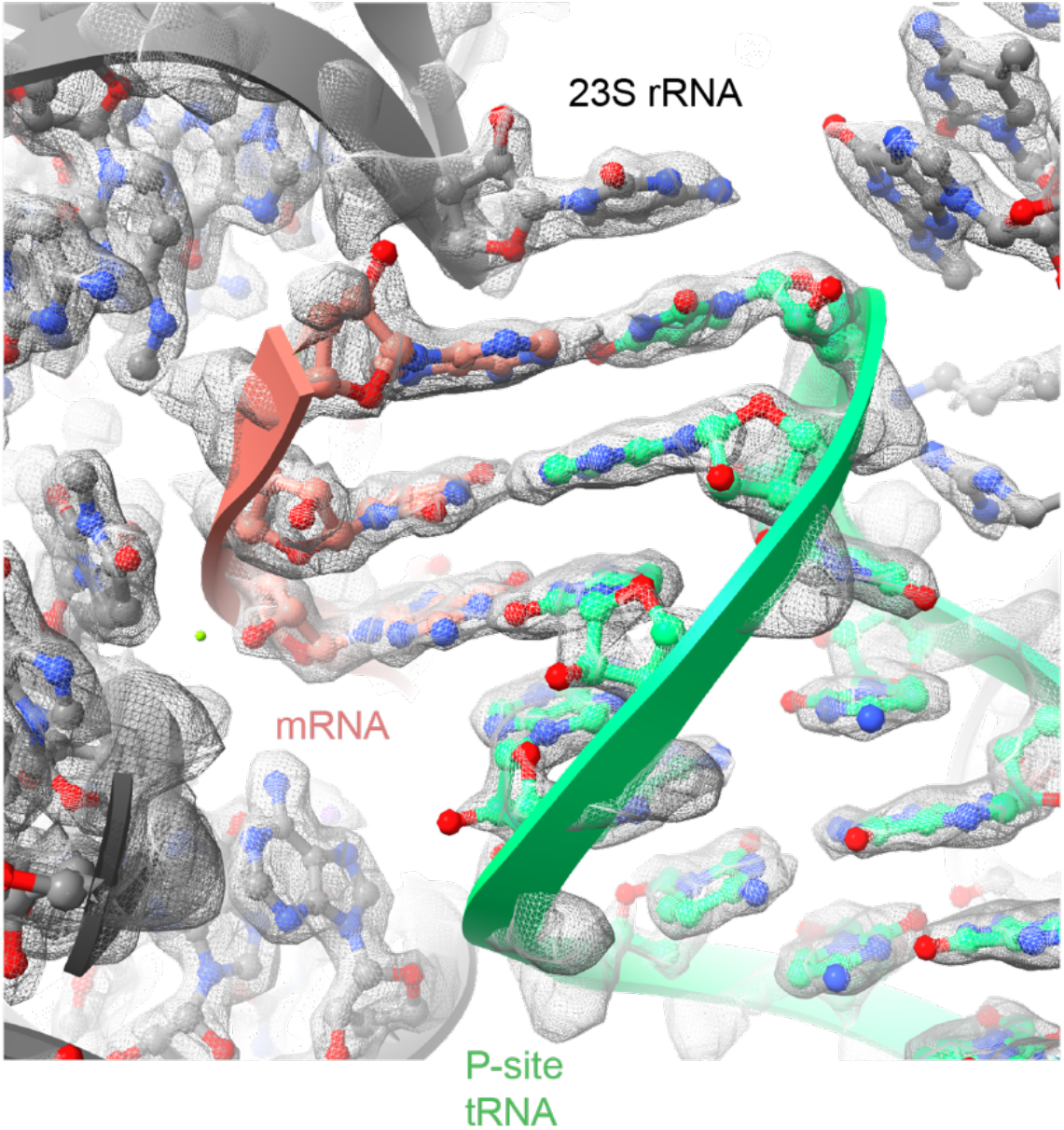
Cryo-EM density corresponding to the codon anti-codon base pairing in the decoding centre of the ribosome. Close-up of the codon-anti codon interactions in the translating ribosome with underlying EM-density shown as mesh. Atomic coordinates are shown as sticks and cartoon illustrations and coloured as in main figure 2.

**Extended Data Fig. 7.**
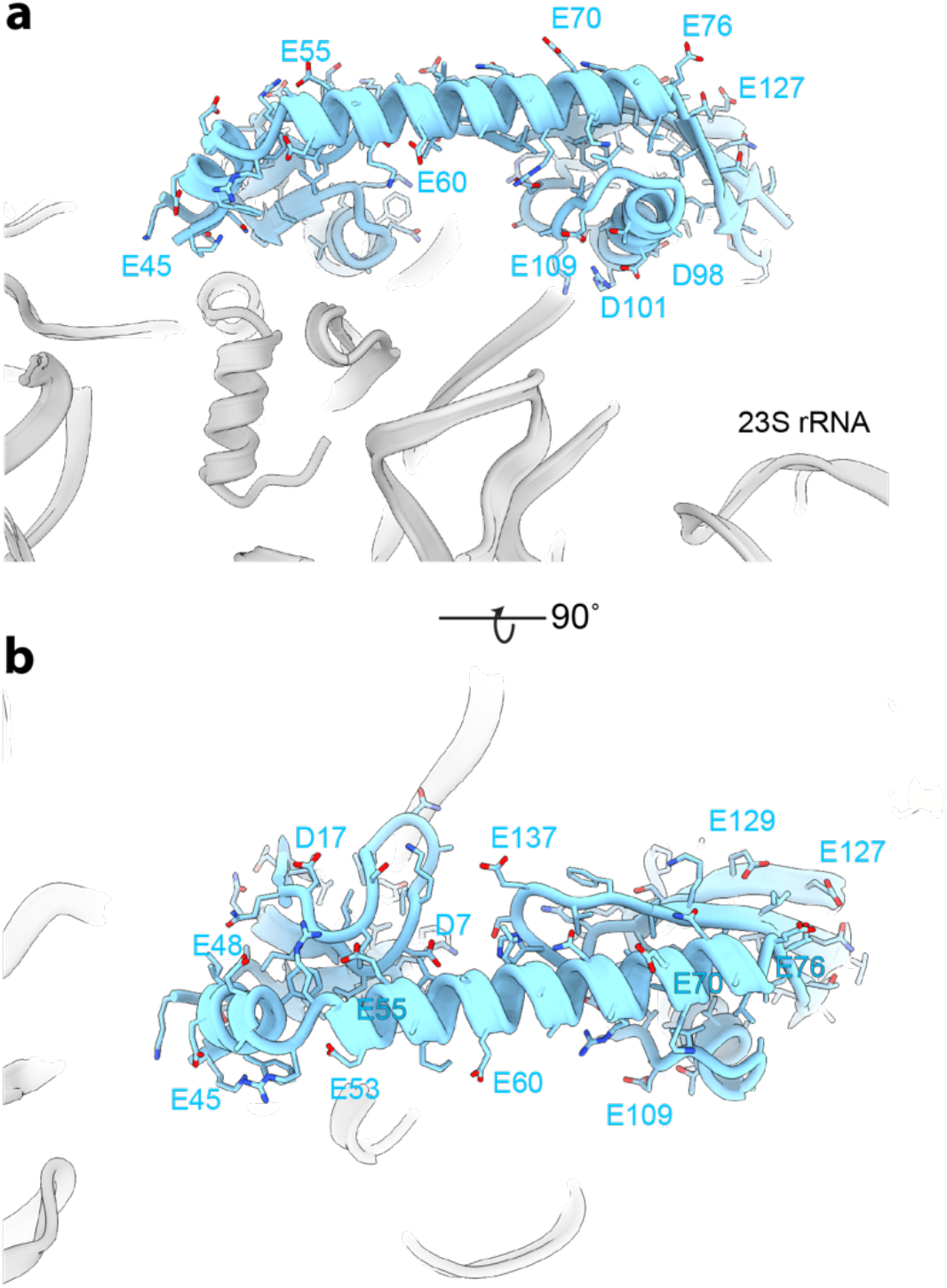
uL9 is rich in negatively charged residues that could repel rRNA. **a** and **b**, Close-ups of the ribosomal protein uL9 shown as a cartoon illustration. Sequence shows high distribution of negatively charged residues. Colouring is the same as in main figures.

**Extended Data Table 1.** Cryo-EM validation and Model Refinements.

|  | 70S ribosome |
| --- | --- |
| EMDB code | #### |
| PDB code | #### |
| **Data collection and processing** |  |
| Magnification | 165,000x (nominal) |
| Voltage (kV) | 300 |
| Electron exposure (e<sup>-</sup>/Å<sup>2</sup>) | 40 |
| Defocus range (μm) | 0.5-1.5 |
| Calibrated Pixel size (Å) | 0.731 |
| Initial particle images (no.) | 1,579,392 |
| Final particle images (no.) | 506,020 |
| Map resolution at FSC=0.143 (Å) | 1.55 |
| **Structure refinement in PHENIX 1.20.1** |  |
| Model resolution at FSC=0.5 (Å) | 1.73 |
| CC<sub>mask</sub> | 0.94 |
| Map sharpening B factor (Å<sup>2</sup>) | - 22.1 |
| **Model composition** |  |
| Non-hydrogen atoms | 151,543 |
| Protein residues | 5,621 |
| RNA residues | 4,448 |
| Ligands: H<sub>2</sub>O/Mg<sup>2+</sup>/K<sup>+</sup> | 12,027 |
| **B factors min/max/mean (Å<sup>2</sup>)** |  |
| Protein | 10.42/120.27/56.09 |
| RNA | 8.70/184.27/49.53 |
| Ligand | 17.89/85.60/40.28 |
| **R.m.s. deviations** |  |
| Bond lengths (Å) | 0.007 |
| Bond angles (°) | 0.703 |
| **Validation** |  |
| MolProbity score | 1.56 |
| Clashscore | 4.12 |
| Poor rotamers (%) | 1.50 |
| **Protein** |  |
| **Ramachandran plot** |  |
| Favored (%) | 96.46 |
| Allowed (%) | 3.34 |
| Disallowed (%) | 0.20 |

**Extended Data Table 2.**
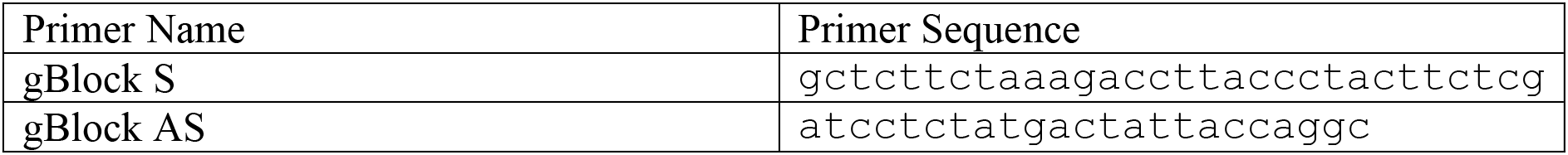
Primers and synthetic DNA sequence for programming ribosomes. Sequence of synthetic DNA used to isolate RNCs: Gctcttctaaagaccttaccctacttctcgccgtaatacgactcactataggcaacaacaacaaccgt tagcttccgacacaaggcttttcactagcaactaaggaggtccaccatggcctacaaagaccacgacg gtgattataaagatcacgacatcgattacaaggacgacgacgacaagtccaaagagccgttgcgcccc cgttgtcgtcctattaatgccacccttgcggtcgagaaagagggctgtccggtttgcatcacggtcaa cactactatttgtgctggatattgccctacggcgacccgtgtccttcagggggtcttgcctgctcttc cccaggttgtatgcaattaccgccgtagtgttactcaccgtatccttactgtccccatcgcgcaggat caagttggcgcgtattatcagcaacctggtcaacaaaacgcaacctggattgttccgccagggaagct gtacaaagctagcgatcaccacctcgggatcagcggcggaaaggtagatatgaacgcctaccaacaac agactaatgtaagaagaaccatgaactgcaaccaggtagatagaaccagcaactttgacaaaatcatc atcgctacagatgctgatggtgatggttagttggggagtaactggttcacatatcactggtctttaca tcggtcttttttacaagcacttcaaagaaatcctgaagaacaacatctacaagcttcagaccccgatt gtcacgctgaaagattctaaaggaaagatcaagaattcccaactgcacgtgcaggtcgacggtgtttt tgcaggacatcatcatcatcatcattaggcaccaccaccaccaccactaggcatcatcatcatcatca ttatgtaccatcgatacgcgttcgaagcttgcggccgcacagctgtatacacgtgcaagccagccaga actcgc

